# DELLA proteins modulate responsive and buffered transcriptional programs in Arabidopsis and rice roots under water limitation

**DOI:** 10.64898/2026.10.22.760938

**Authors:** Cynthia Wong, David Alabadí, Miguel A. Blázquez, Bruno Catarino

## Abstract

Water limitation constrains plant growth and productivity, yet the molecular mechanisms regulating root responses remain incompletely understood. The DELLA repressors of gibberellin signaling are key growth regulators under stress, but their role in shaping transcriptional responses to water limitation, and its conservation, remain unclear. Here, we investigated DELLA function in roots of *Arabidopsis thaliana* and *Oryza sativa* under reduced water availability simulated by polyethylene glycol (PEG). Loss of DELLA function reduced the sensitivity of root growth to water limitation, affecting primary root elongation in *A. thaliana* and crown root formation in *O. sativa*. Transcriptomic profiling revealed extensive DELLA-dependent regulation, partitioned into two modes: (i) genes requiring DELLAs for appropriate responses to water limitation (“WT –responsive”), and (ii) genes whose responsiveness is detected predominantly in the absence of DELLAs (“buffered”, an operational label for an expression pattern, not a demonstrated mechanism). These patterns suggest that DELLAs both enable specific responses and apparently constrain broader transcriptional responsiveness under osmotic stress. Although the requirement for DELLA function is conserved, the developmental outputs and downstream transcriptional programs are largely species-specific. Together, our results identify DELLAs as key modulators of root responses to water limitation, with a proposed dual role in shaping and limiting transcriptional dynamics.

**Highlight:** DELLAs coordinate root adaptation to reduced water availability by enabling selective stress responses while apparently constraining broader transcriptional programs in angiosperms.

## Introduction

Plants have evolved sophisticated, interconnected signaling pathways to perceive and translate environmental stimuli into adaptive growth and developmental responses (Lamers *et al.,* 2020). Among these stimuli, abiotic stresses, particularly drought, exert a profound impact on plant fitness and survival (Gupta *et al*., 2020; Gampe *et al.,* 2021; Munns and Millar, 2023; Guo *et al*., 2026). As global climate models predict intensifying drought cycles throughout the twenty-first century, the resulting instability poses a dire threat to agricultural productivity and ecosystem resilience (Sillmann *et al.,* 2013; Lehmann *et al.,* 2015; Yin *et al*., 2018). Recent decades have already seen drought-induced losses of approximately 10% in major crops like maize and soybean; by 2050, productivity in key staples such as rice is projected to decline by 20% or more in high-risk regions (Kraklow *et al.,* 2025). Consequently, because roots directly perceive changes in soil water potential and initiate adaptive responses, elucidating the molecular mechanisms governing root-specific responses to water limitation is a critical priority for ensuring global food security.

Water limitation triggers extensive morphological and physiological adjustments in the rooting system (Shoaib *et al.,* 2022; Voothuluru *et al.,* 2024). Plants dynamically modulate root architecture to optimize water foraging, including changes in primary root elongation, lateral root formation, and, in monocots, the initiation of crown roots (Sebastian *et al.,* 2016; Ranjan *et al.,* 2022). While mild water deficits can promote deeper rooting, more severe stress conditions generally restrict growth to maintain cellular homeostasis (de Ollas *et al*., 2023). Importantly, transcriptomic studies have demonstrated that root responses to water limitation are not simply scaled versions of shoot responses, but instead involve distinct temporal dynamics and pathway activation, often with earlier and organ-specific transcriptional remodeling (Minh –Thu *et al.,* 2013; Rasheed *et al.,* 2016; Royer *et al.,* 2016; Gupta *et al.,* 2020).

A major hormonal pathway implicated in these responses is the gibberellin (GA) signaling pathway. A large body of evidence indicates that water limitation is frequently associated with reduced GA accumulation or responsiveness, resulting in the stabilization of DELLA proteins, which act as key repressors of GA-mediated growth (Claeys and Inzé, 2013; Liao *et al*., 2023). This molecular shift is commonly linked to growth and enhanced stress tolerance. However, this relationship is not universal and depends on developmental context, organ, and experimental conditions, as well as on the physiological or agronomic traits being measured (Claeys *et al.,* 2012; Dubois *et al*., 2013; Lo *et al.,* 2017; Wang *et al.,* 2020, 2021; Liao *et al.,* 2023; Swift *et al.,* 2026). Thus, rather than representing a simple linear pathway, GA-DELLA signaling operates as a context-dependent regulatory module that integrates environmental and endogenous cues.

DELLA proteins are transcriptional regulators belonging to the GRAS family and act as conserved central hubs within these networks (Hernández –García *et al.,* 2021; Briones-Moreno *et al.,* 2023). They modulate gene expression through interactions with numerous transcription factors (TFs), influencing diverse biological processes including cell proliferation, differentiation, and stress responses (Alabadí and Sun, 2025). Through mechanisms such as the physical sequestration of TFs, interference with chromatin-bound complexes, and recruitment of co-regulators, DELLAs coordinate growth regulation with environmental acclimation (Alabadí and Sun, 2025). This positions DELLAs as nodes that balance developmental programs with stress-responsive signaling pathways (Colebrook *et al.,* 2014).

Despite substantial progress in understanding GA-DELLA signaling under water limitation, most causal evidence derives from shoots, leaves, or whole plant analyses. In contrast, the role of DELLAs in root-specific responses remains relatively underexplored. Transcriptomic studies in *Arabidopsis thaliana* and *Oryza sativa* have shown that roots deploy distinct genotype –and time –dependent components (Minh-Thu *et al*., 2013; Baldoni *et al*., 2016; Rasheed *et al.,* 2016; Royer *et al*., 2016; Todaka *et al.,* 2017). However, these datasets rarely combine organ-resolved transcriptomics with direct genetic perturbations of GA signaling or DELLA function, leaving a gap in our understanding of how these regulators shape root-specific transcriptional responses.

An additional layer of complexity arises from the diversity of experimental systems used to impose water limitation. Soil drying, dehydration, and osmotic treatments such as polyethylene glycol (PEG) are often used interchangeably, yet they do not produce equivalent physiological or transcriptional responses (Gonzalez *et al.,* 2024). Recent studies demonstrate that PEG-based and other low water potential assays recapitulate only part of the drought response and can diverge substantially from soil-based drought at the transcriptomic level (Kylyshbayeva *et al.,* 2025). Moreover, early root responses depend on the physical nature of the stress signal, including the relative contributions of osmotic and turgor components (Crabos *et al.,* 2023). Complementary physiological work further shows that PEG induces dose-dependent root responses governed by specific hormonal and hydraulic mechanisms (Rosales *et al.,* 2019). Therefore, to ensure high-resolution spatiotemporal control over water potential, minimizing the confounding variables inherent in heterogeneous soil drying, we utilized a defined PEG-induced osmotic gradient. This approach allowed us to isolate the DELLA-dependent components of the hydraulic and osmotic response from secondary stress signals.

To address these gaps, we conducted comparative phenotypic and transcriptomic analyses of *A. thaliana* and *O. sativa* roots exposed to PEG –induced water limitation. By combining organ-resolved transcriptomics with genetic perturbation of DELLA function, we investigated how these conserved regulators shape root-specific responses and to what extent their downstream transcriptional networks are conserved or have diverged between these two angiosperm lineages. Our findings reveal that DELLAs mediate root responses to PEG-induced water deficit through a dual mode of action: enabling specific transcriptional responses while apparently constraining others, a pattern consistent with, although not demonstrating, a transcriptional buffering role.

## Materials and methods

### Plant material and growth conditions

*Arabidopsis thaliana* ecotype Landsberg *erecta* (L*er*) and the loss-of-function mutant At*dellaKO* (L*er*) (Feng *et al*., 2008) were used. Seeds were surface –sterilized in 70% (v/v) ethanol for 15 min, followed by 100% ethanol for 2 min under gentle agitation, then dried in a laminar flow hood. Seeds were plated individually (ca. 0.5 cm spacing) on half-strength Murashige and Skoog (MS) medium (pH 5.7), supplemented with 0.8% phytoagar in square Petri dishes. After stratification at 4°C for four days in darkness, seedlings were grown vertically under long-day conditions (16 h light/8 h dark; 30 µmol m^-2^ s^-1^) at 22°C. Five days after germination, 12 seedlings per plate were transferred to PEG-infused plates (van der Weele *et al*., 2000) and grown vertically for additional 6 days.

*Oryza sativa* cv. Nipponbare (WT) and the loss-of-function mutant Os*slr1-1* (Ikeda *et al*., 2001) (henceforth referred to as Os*slr1*) were used. Seeds were manually dehusked and surface-sterilized with 70% (v/v) ethanol for 30 s, followed by 60% (v/v) bleach for 30 min, then rinsed six times with sterile water. Seeds were imbibed for ca. 2 h and plated on 0.8% phytoagar prepared in tap water (pH 5.7). Ten germinated seeds per plate were transferred to PEG-infused plates and grown vertically for 3 days under continuous white light (100 µmol m^-2^ s^-1^) at 28°C prior to phenotypic analyses and tissue collection. Given the sterility of Os*slr1* mutants, the seed stock is maintained as a segregating population. Homozygous mutant seedlings were identified for each experiment through genotyping, followed by Sanger sequencing, of the mutated locus using the following oligos: Fwd –GTGCCGGTTGTGGTGGTTGA and Rev –GAGTCCCCGGTACTGGAAGT.

### Preparation of PEG-infused plates via the overlay method

PEG-infused plates were prepared using the overlay method (van der Weele *et al*., 2000). PEG8000 (Sigma-Aldrich; CAT #: P5413-500G) solutions (0-30%, w/v) were prepared in half-strength MS medium (for *A. thaliana*) or tap water (for *O. sativa*), adjusted to pH 5.7, sterilized by vacuum filtration, and overlaid onto solidified agar medium at 1:1 volume ratio. Plates were incubated for 24 h to allow equilibration of water potential, after which excess liquid was removed. Mock plates (0% PEG) were prepared using the corresponding liquid medium without PEG8000. For *O. sativa*, a portion of the agar surface was removed to facilitate stable placement of seedlings during vertical growth.

### Root phenotype quantifications

For *A. thaliana*, primary root tips were marked at the time of transfer to PEG plates (day 0) and every two days thereafter. Plates were imaged at day 6, and root growth was quantified using Fiji (ImageJ) (Schindelin *et al*., 2012; Schneider *et al*., 2012). Growth rates were calculated by measuring root elongation between successive time points. For *O. sativa*, primary root length was measured over a 3-day treatment period using the same approach. In addition, the number of crown roots per seedling was manually counted at the end of the treatment. All measurements were calibrated using a scale included in the images. Crown root numbers were treated as count data and analyzed using a generalized linear model (quasipoisson family) followed by post-hoc comparisons with estimated marginal means (‘emmeans’ package in R). Sample sizes varied among *O. sativa* genotype–concentration combinations, primarily reflecting the recovery of homozygous Os*slr1* individuals from segregating populations, together with differences in germination and seedling recovery among treatments; the exact n for every genotype and concentration is indicated in Fig. 1 and Supplementary Fig. S1. All raw measurements, statistical analyses and representative images of *A. thaliana* seedlings are provided in Supplementary Data S1.

**Figure 1.**
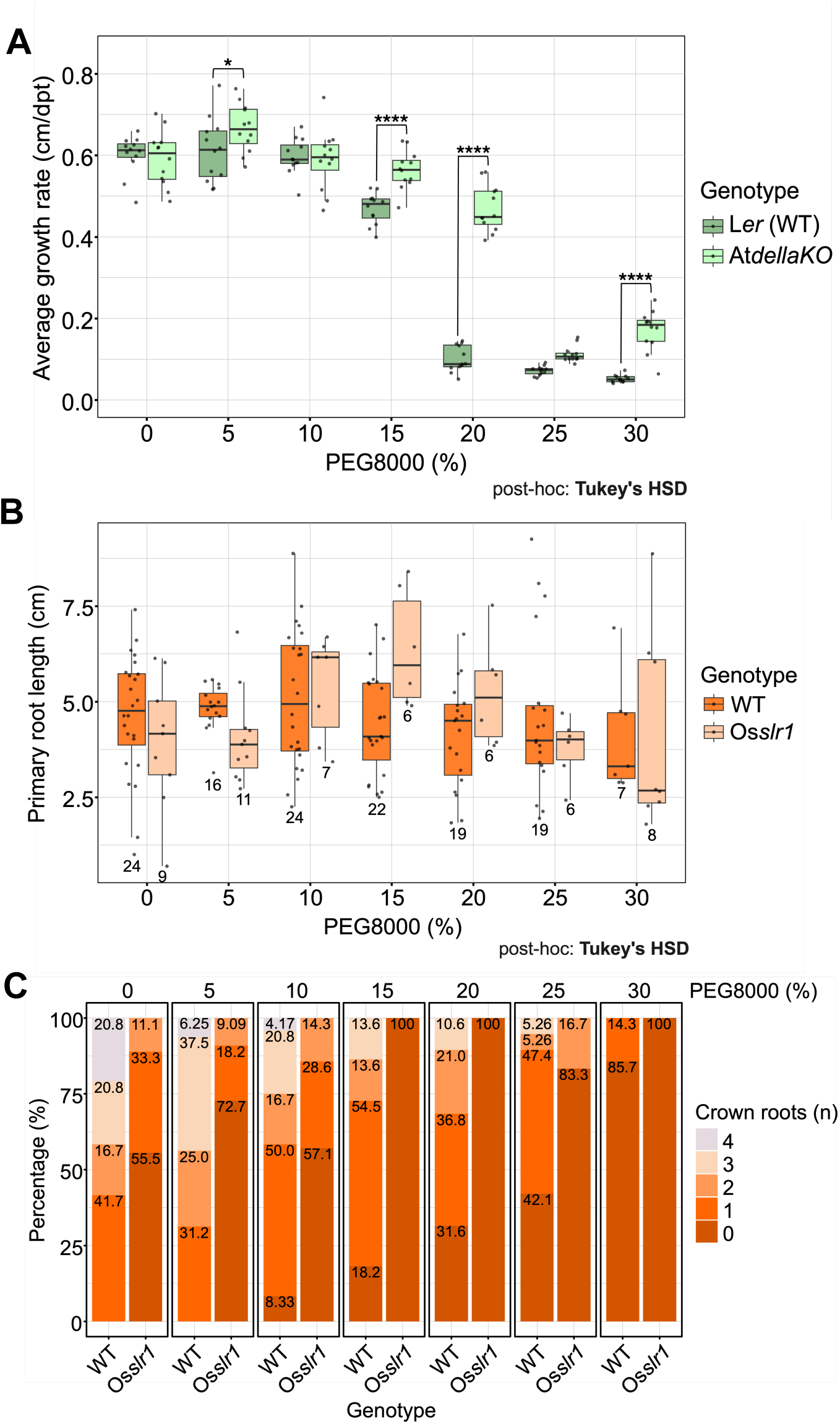
DELLA proteins modulate root growth responses to PEG-induced reduction of water potential in *A. thaliana* and *O. sativa*. **A)** Primary root growth rates of *A. thaliana* Ler (WT) and At*dellaKO* seedlings (n = 12 per condition and genotype) across a gradient of PEG8000 concentrations (0-30%). **B)** Primary root length of *O. sativa* Nipponbare (WT) and Os*slr1* across the same PEG gradient; no significant differences between genotypes were detected after correction for multiple comparisons. Numbers below boxplots indicate sample size (n) per condition and genotype. **C)** Number of crown roots in Nipponbare (WT) and Os*slr1* across increasing PEG concentrations, shown as the percentage of seedlings per crown-root-number category; individual counts and statistics are provided in Supplementary Fig. S1. Statistical significance in A) and B) was determined by two-way factorial ANOVA followed by Tukey’s HSD post-hoc tests, and crown root numbers were analyzed using a generalized linear model (quasipoisson) followed by emmeans post-hoc comparisons (Supplementary Fig. S1). Asterisks indicate significant differences between WT and mutant genotypes at a given PEG concentration (* p < 0.05; **** p < 0.0001). Raw measurements and complete statistics are provided in Supplementary Data S1. **Alt text:** Box plots comparing primary root growth rate in *A. thaliana* and primary root length in *O. sativa* between wild type and DELLA mutants across seven PEG concentrations, plus stacked bars of *O. sativa* crown root number categories.

### RNA extraction

Whole roots were harvested from independent biological replicates (three per condition and genotype). The number of roots pooled per replicate was 12 for *A. thaliana* and 4 for *O. sativa*. Samples were flash-frozen in liquid nitrogen and stored at –80°C. Total RNA was extracted using the NucleoSpin RNA Plant kit (Macherey-Nagel) following the manufacturer’s instructions. RNA quantity, quality, and integrity were assessed using Agilent Bioanalyzer or Fragment Analyzer systems prior to sequencing.

### RNA sequencing

RNA-seq libraries were prepared from three biological replicates per genotype and condition (mock and 25% PEG). The 25% PEG8000 concentration was selected because it imposed a severe but sub-lethal stress in both species while preserving sufficient root material for RNA extraction; although primary root growth rates of the two *A. thaliana* genotypes did not differ significantly at this concentration, the treatment elicited clear DELLA-dependent transcriptional changes (see Results). We note that comparable root elongation does not necessarily imply an equivalent developmental stage (see Discussion). Sequencing was performed by BGI using the DNBSeq^TM^ platform with paired –end 100 bp reads, generating over 24 million reads per sample. After adapter removal and quality trimming of raw reads with cutadapt (Martin, 2011), a quality analysis of clean reads was done with FastQC (https://www.bioinformatics.babraham.ac.uk/projects/fastqc/). Clean read pairs longer than 20 nt were then mapped to the *A. thaliana* reference genome (TAIR10) (Lamesch *et al*., 2012) and the *O. sativa* IRGSP-1.0 reference genome (Kawahara *et al*., 2013; Sakai *et al*., 2013) using STAR (Dobin *et al*., 2013). The number of read counts mapped to one and only one of the annotated genes in the genome (uniquely mapped) was obtained using STAR and samtools (Danecek *et al*., 2021), with the Araport11 (Cheng *et al*., 2017) and the IRGSP-1.0 (Kawahara et al., 2013; Sakai *et al*., 2013) structural annotations, for *A. thaliana* and *O. sativa*, respectively.

#### Differential gene expression analysis

Differential expression analysis was performed using DESeq2 (Love *et al*., 2014). Genes were considered differentially expressed with an adjusted p –value < 0.01 and absolute log_2_ fold change ≥ 1. Genotype-by-environment (G×E) interaction effects were assessed using the DESeq2 model including genotype, treatment, and their interaction. The complete differential expression and G×E outputs are provided in Supplementary Data S2. The numbers of up –and downregulated genes in each differentially expressed gene set are provided in Supplementary Data S3. Reproducibility across biological replicates was assessed by principal component analysis of normalized expression values from all libraries (Supplementary Fig. S2).

#### Hierarchical clustering

Normalized expression values of transcripts per million (TPM) were transformed into Z-scores across samples. Hierarchical clustering was performed with ‘hclust’ function from the R ‘stats’ package (v 4.4.1.) using Pearson correlation-based distance and complete linkage. The optimal number of clusters was determined using silhouette analysis, and genes were grouped into 20 clusters. The optimal number of clusters was established using Silhouette analysis for a range of *k* = 2 to 30. Based on the average silhouette width, the dendrogram was partitioned into 20 discrete clusters using the ‘cutree’ function (Supplementary Data S4). The 20 clusters per species were visualized using the R package ‘ggplot2’ (Wickham, 2016).

#### GxE heatmap visualization

Genes showing significant GxE interaction effects (p < 0,05) and consistent differential expression (p < 0.01) were selected and visualized using Morpheus (Broad Institute; https://software.broadinstitute.org/morpheus). Genes were classified based on their response patterns into WT-responsive or buffered categories (Supplementary Data S5).

#### Two-dimensional density plots

Density plots comparing log_2_ fold changes between WT and mutant genotypes were generated using ‘ggplot2’ (Wickham, 2016) on R (v. 4.4.1). PEG-responsive genes (p < 0.01) were included, and genes classified as buffered based on GxE analysis were highlighted.

#### Gene Ontology enrichment analysis

GO enrichment analysis was performed using ShinyGO 0.85 (Ge *et al*., 2020). Redundant GO terms were summarized using REVIGO (Supek *et al*., 2011), using the SimRel semantic-similarity measure and the default redundancy cutoff (0.7), and enriched terms were grouped into broader functional categories for visualization. For the cross-species comparison, the REVIGO network was exported to Cytoscape (Shannon et al., 2003) for visualization; edges represent semantic similarity between GO terms rather than biological interactions, and no additional edge-weight threshold was applied in Cytoscape (edge weights ranged from 0.278 to 0.700). Dot plots were generated using ‘ggplot2’ (Wickham, 2016) on R (v. 4.4.1) for each species. Enriched GO terms, their full names and the assigned functional categories are listed in Supplementary Data S6. An equivalent GO enrichment analysis was performed on the orthologous DELLA-dependent genes identified in the cross –species comparison (see Results).

#### Cross-species comparison of orthologous genes

Orthologous relationships between *A. thaliana* and *O. sativa* genes were obtained from Ensembl Plants (release 62) (Howe *et al*., 2020; Martin *et al*., 2023). Differential expression patterns of conserved DEGs were compared across species and visualized using heatmaps generated on Morpheus (Broad Institute; https://software.broadinstitute.org/morpheus).

Ortholog relationships between the two species are frequently many-to –many; where one species contributes more genes to an orthogroup than the other, the surplus positions have no counterpart in the other species and are shown in grey in the cross-species heatmap. Rows of the heatmap were ordered manually to group orthogroups with related expression patterns. To quantify cross-species concordance, the ratio between the *O. sativa* and *A. thaliana* log_2_ fold-change values was computed for each orthologous gene pair in each of the four comparisons (PEG/mock in each genotype, and mutant/wild type in each condition); orthogroups with positive ratios in all comparisons were classified as responding in the same direction, orthogroups with a negative ratio in at least one comparison as responding in opposite directions, and orthogroups whose member pairs behaved inconsistently as discrepant.

#### Transcription factor association studies

Putative transcriptional associations were inferred based on the enrichment of TF binding sites in promoter regions (–500 bp to +100 bp from TSS) using the PlantRegMap “Regulation Prediction” tool (Tian *et al*., 2020). TFs associated with DELLA-dependent gene sets were identified separately from WT-responsive and buffered genes as indicated in the corresponding figures. To facilitate visualization, TF-TF relationships were represented in Cytoscape (Shannon *et al*., 2003). For clarity, the displayed networks were restricted to a subset of nodes most relevant to DELLA-associated regulation. These included: (i) TFs previously reported to interact with DELLAs in *A. thaliana* (la Rosa *et al*., 2014; Lantzouni *et al*., 2020), and (ii) TFs that are differentially expressed in response to PEG treatment and are directly connected to DELLA-interacting TFs based on predicted associations. Comparative analyses between *A. thaliana* and *O. sativa* were performed by assessing binding-site enrichment of orthologous TFs in each species. Importantly, these analyses represent inferred associations mostly based on motif enrichment and do not establish direct regulatory interactions.

#### Analysis of conserved TF targets

Differential expressions of predicted target genes of conserved TFs were analyzed using a Linear Mixed-Effects Model (‘lme4’ package in R (v. 4.4.1)). Species and treatment were included as fixed effects. Post-hoc comparisons were performed using estimated marginal means (‘emmeans’ package). Data were visualized using violin plots generated using the log_2_ fold change values of target genes with ‘ggplot2’ (Wickham, 2016) on R (v. 4.4.1).

## Results

### DELLAs mediate root growth responses to PEG-induced water limitation in *A. thaliana* and *O. sativa*

To examine root responses to PEG-induced water limitation, we performed a gradient analysis using increasing concentrations of PEG8000 (0–30%) in *A. thaliana* and *O. sativa*. In *A. thaliana*, primary root growth rates were measured in the wild type (L*er*) and in the At*dellaKO* quintuple mutant (Feng *et al.,* 2008). Under control conditions (0% PEG8000), both genotypes exhibited similar growth rates (Fig. 1A). Increasing PEG concentrations progressively reduced growth rates in both genotypes; however, this reduction was more pronounced in the wild type than in *AtdellaKO*, with significant differences between genotypes at 5%, 15%, 20% and 30% PEG8000 (Fig. 1A; Supplementary Data S1). The dose–response was not strictly monotonic in At*dellaKO* (e.g. at 25% PEG8000 the genotype difference was not significant after correction for multiple comparisons); we therefore interpret the response to the gradient as a whole rather than emphasizing any individual concentration. These results indicate that DELLA function contributes to the sensitivity of root growth to PEG –induced water limitation in *A. thaliana* roots.

We performed a similar phenotypic analysis in *O. sativa*. Seedlings were grown in the same PEG8000 gradient, and root growth was assessed in the wild type (Nipponbare) and in the Os*slr1-1* loss-of-function mutants (Ikeda *et al.,* 2001). Because Os*slr1* mutants are sterile, F1 progeny from heterozygous plants were genotyped to identify homozygous mutant individuals. In contrast to *A. thaliana*, primary root growth in *O. sativa* was not strongly affected across the PEG gradient in either genotype (Fig. 1B). No significant differences between genotypes were detected at any PEG concentration after correction for multiple comparisons (Fig. 1B; Supplementary Data S1); we therefore do not interpret primary root length as an OsSLR1-dependent trait under these conditions. Interestingly, the distribution of crown root numbers in wild type plants progressively decreased with increasing PEG concentrations, indicating that PEG-induced water limitation negatively regulates crown root initiation in *O. sativa* (Fig. 1C; Supplementary Fig. S1). *Osslr1* mutants exhibited significantly fewer crown roots than the wild type under mock conditions and at 5% and 10% PEG8000 (quasipoisson generalized linear model; Supplementary Fig. S1; Supplementary Data S1), but notably lacked the dose-dependent shift observed in wild type (Fig. 1C). This reduced responsiveness suggests that OsSLR1 is required for the modulation of crown root development in response to PEG –induced water limitation. Together, these results indicate that DELLA proteins contribute to the regulation of root growth and architecture under PEG-induced water limitation in both *A. thaliana* and *O. sativa*, although the specific developmental outputs differ between species.

### The root transcriptional responses to PEG-induced water limitation are regulated by AtDELLAs in *A. thaliana*

To investigate the role of AtDELLAs in transcriptional responses to PEG –induced water limitation, we performed RNA-seq on roots of wild type and At*dellaKO* seedlings grown under mock condition and 25% PEG8000. Principal component analysis of all biological replicates showed that samples grouped according to treatment and genotype, supporting the reproducibility of the dataset (Supplementary Fig. S2A). Differentially expressed genes (DEGs) were identified using a fold change of ± log_2_(2) and adjusted p-value < 0.01. Comparison between PEG-treated and control samples revealed 833 genes that were differentially expressed exclusively in the wild type and 950 genes exclusively in At*dellaKO* (Fig. 2A; Supplementary Data S2). Among the WT –exclusive genes, 258 were upregulated and 575 downregulated, whereas 346 of the At*dellaKO*-exclusive genes were upregulated and 604 downregulated; the corresponding breakdown for all gene sets is provided in Supplementary Data S3. This distribution indicates that AtDELLAs modulate the transcriptional response to PEG-induced water limitation, both by enabling specific responses in the wild type, and by restricting responses that become apparent in their absence. To characterize transcriptional behaviors more broadly, hierarchical clustering of DEGs identified 20 clusters showing distinct expression patterns (Fig. 2B; Supplementary Data S4). Most clusters exhibited altered PEG responsiveness in At*dellaKO* compared to wild type (Fig. 2B), indicating that AtDELLAs contribute to shaping transcriptional responses under these conditions.

**Figure 2.**
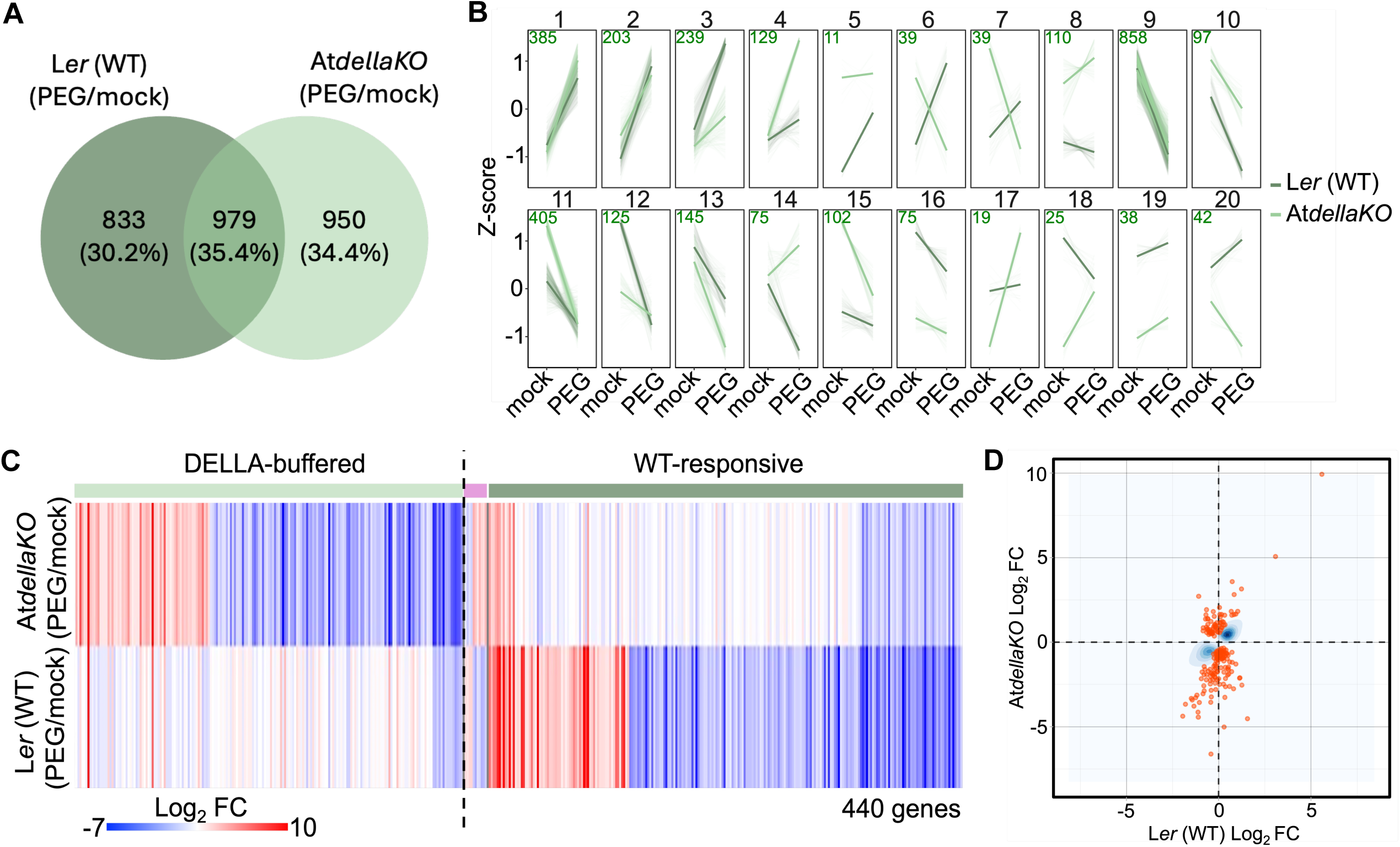
DELLA-dependent transcriptional responses to PEG-induced water limitation in *A. thaliana* roots. **A)** Venn diagram illustrating the overlap of genes differentially expressed in response to PEG treatment (PEG vs mock) in wild type (WT) and At*dellaKO* seedlings. Number of genes responding exclusively in WT or in the mutant are indicated. **B)** Hierarchical clustering of differentially expressed genes (DEGs) across WT and At*dellaKO* samples under mock and PEG conditions, identifying 20 clusters with distinct expression patterns. **C)** Heatmap showing significant genotype-by-environment (GxE) interaction effects (p < 0.05), highlighting two categories: genes responsive to PEG predominantly in WT (“WT-responsive”) and genes whose responsiveness is more evident in the mutant background (“buffered”). **D)** Two-dimensional density plot comparing log_2_ fold changes of PEG-responsive genes (p < 0.01) in WT and At*dellaKO*. Genes classified as “buffered” based on GxE analysis are highlighted. “Buffered” is used throughout as an operational label for genes whose PEG response is detected predominantly in the mutant background; it does not imply a demonstrated buffering mechanism. **Alt text:** Venn diagram of PEG-responsive genes in wild type versus At*dellaKO*, line plots of twenty expression clusters, a heatmap of genotype-by-environment genes split into WT-responsive and buffered groups, and a density scatter plot of fold changes.

We next classified the DELLA-dependent gene expression into two categories. The first one, termed “AtDELLA-dependent WT-responsive”, consists of genes whose differential expression between PEG and mock conditions is observed in the wild type but altered or absent in At*dellaKO* (Fig. 2A and Fig. S3). The second category, termed “AtDELLA-dependent buffered”, comprises genes that respond to PEG only (or primarily) in the mutant background, suggesting that their responsiveness is reduced or constrained in the presence of DELLA proteins (Fig. 2A). Throughout this study, “buffered” is used as an operational label for this expression pattern and does not imply a demonstrated buffering mechanism; such a pattern is consistent with direct restraint by DELLAs, but could also arise from differences between genotypes in basal expression, developmental stage, root cellular composition or perceived stress severity, possibilities that our whole-root experimental design cannot distinguish (see Discussion). Employing Gene-by-Environment Interaction (GxE) statistical analyses on our transcriptomes, we further supported the occurrence of the DELLA-mediated modes of response mentioned above. 56.1% of the GxE DEGs are AtDELLA-dependent WT –responsive, whereas 43.9% of the GxE DEGs are AtDELLA-dependent buffered genes (Fig. 2C; Supplementary Data S5). Furthermore, most of the AtDELLA –dependent buffered DEGs, based on the GxE statistical analysis, overlap with the DEGs identified above which were obtained using the comparative analyses between our transcriptomes, and the majority of these is downregulated by PEG in the At*dellaKO* mutant (Fig. 2C and D). These observations are consistent with AtDELLAs dampening the transcriptional response to PEG-induced water limitation in a subset of genes.

Gene Ontology (GO) analysis revealed both shared and distinct biological processes associated with these categories (Fig. 3; Supplementary Data S6). Processes such as nitrogen metabolism, response to oxygen levels, and hormone signaling were enriched in both categories, while developmental processes and light signaling-related pathways were preferentially associated with the buffered category (Fig. 3; Supplementary Data S6). These results indicate that AtDELLAs both modulate and appear to constrain overlapping and distinct transcriptional responses to PEG-induced water limitation.

**Figure 3.**
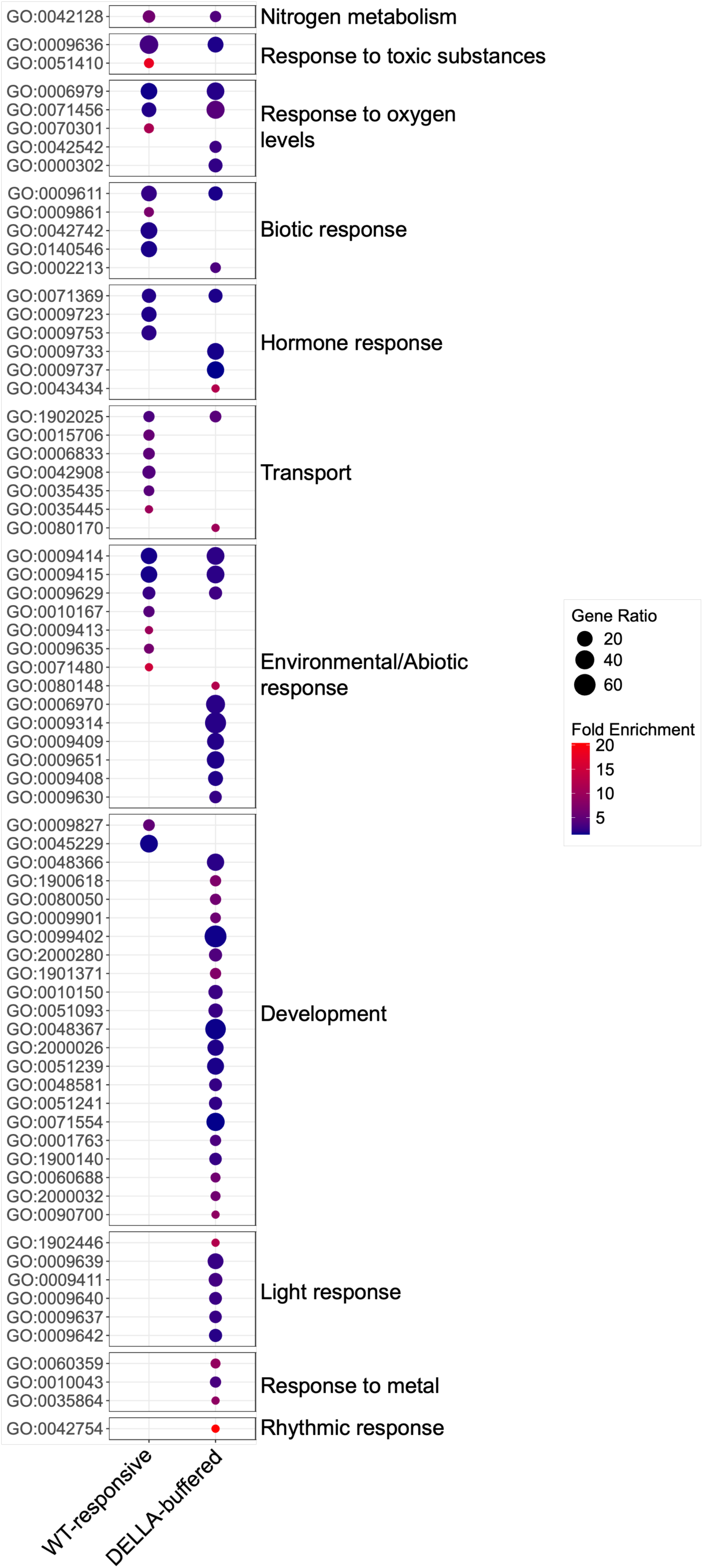
Functional enrichment of DELLA-dependent gene expression categories in *A. thaliana* roots. Dot plot showing Gene Ontology (GO) biological processes enriched among PEG-responsive DEGs classified as WT-responsive or buffered. Dot size represents the number of genes associated with each term, and color intensity indicates the degree of enrichment. Shared and distinct functional categories are observed between the two gene sets. Labels on the right correspond to author-assigned functional groupings of the non-redundant GO terms obtained after REVIGO analysis, not to parent GO terms; full GO identifiers and term names are listed in Supplementary Data S6. **Alt text:** Dot plot of enriched Gene Ontology biological processes for WT –responsive and buffered gene sets in *A. thaliana*, with dot size showing gene number and colour showing enrichment, grouped into functional categories.

To further explore the regulatory basis of these responses, we inferred the TFs with enriched binding sites in each gene set (see Materials and Methods for detailed criteria). While the buffered network displayed a more complex structure, eight TFs were shared between the WT-responsive and buffered networks, suggesting a partial overlap in regulatory components (Fig. S4; Supplementary Data S7). Overall, these analyses suggest that AtDELLAs regulate root transcriptional responses to water limitation by modulating specific gene sets and acting as a broader buffer on transcriptional responsiveness.

### OsSLR1 modulates root transcriptional responses to PEG-induced water limitation in *O. sativa*

To determine the role of OsSLR1 in *O. sativa* root transcriptional responses, we performed RNA-seq on wild type and *Osslr1* mutant roots under mock condition and 25% PEG8000. Homozygous mutant seedlings were identified by genotyping prior to RNA extraction. Differential expression analysis revealed 191 genes that were PEG-responsive exclusively in the wild type and 388 genes exclusively in the *Osslr1* mutants (Fig. 4A; Supplementary Data S2). Among the WT-exclusive genes, 88 were upregulated and 103 downregulated, whereas 35 of the Os*slr1*-exclusive genes were upregulated and 353 downregulated (Supplementary Data S3). Principal component analysis of all *O. sativa* replicates likewise separated samples by genotype and treatment (Supplementary Fig. S2B). This distribution indicates that OsSLR1 contributes to shaping the transcriptional response to PEG-induced water limitation in *O. sativa* roots.

**Figure 4.**
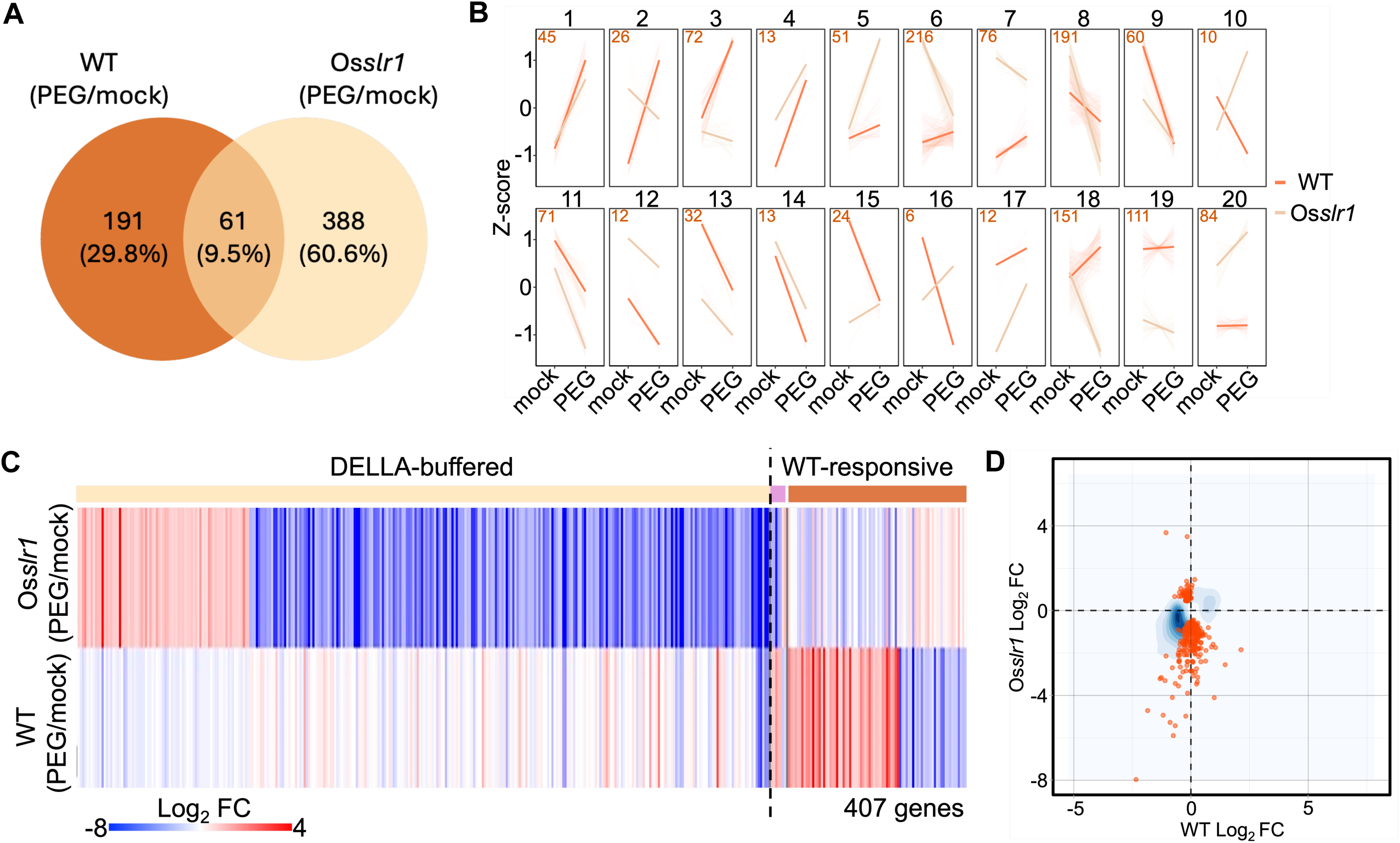
OsSLR1-dependent transcriptional responses to PEG-induced water limitation in *O. sativa* roots. **A)** Venn diagram showing the overlap of PEG –responsive genes in wild type (WT) and Os*slr1* seedlings. **B)** Hierarchical clustering of DEGs across WT and Os*slr1* samples, identifying 20 clusters with distinct expression profiles. **C)** Heatmap of GxE DEGs (p < 0.05), showing genes responsive predominantly in WT (“WT-responsive”) and genes whose responsiveness is more evident in the mutant (“buffered”). **D)** Two-dimensional density plot comparing log_2_ fold changes in WT and Os*slr1*, with buffered genes highlighted. “Buffered” is used as an operational label as defined in Fig. 2. **Alt text:** Venn diagram of PEG-responsive genes in rice wild type versus Os*slr1*, line plots of twenty expression clusters, a heatmap of genotype-by-environment genes split into WT-responsive and buffered groups, and a density scatter plot of fold changes.

Hierarchical clustering analysis identified 20 gene clusters with distinct expression profiles, most of which exhibited altered PEG responsiveness in the mutant background, consistent with OsSLR1-dependent transcriptional regulation (Fig. 4B; Supplementary Data S4). As in *A. thaliana*, we classified OsSLR1-dependent gene expression into WT-responsive and buffered categories (Fig. 4A and Fig. S3). Statistical analysis using GxE on our four *O. sativa* transcriptomes, supported the evidence that DELLA-mediated gene expression encompasses the two modes of response mentioned above, defined operationally as in *A. thaliana*. 21.9% of the GxE DEGs are OsSLR1-dependent WT-responsive, whereas 78.1% of the GxE DEGs are OsSLR1-dependent buffered genes (Fig. 4C; Supplementary Data S5). Moreover, a great proportion of the OsSLR1-dependent buffered genes, based on the GxE statistical analysis, overlap with the DEGs identified through the comparative analyses between our *O. sativa* transcriptomes (Fig. 4D). Interestingly, most of these OsSLR1-dependent buffered genes are downregulated by PEG in the Os*slr1* loss-of-function mutant. (Fig. 4C and D). This is consistent with a role for OsSLR1 in limiting or constraining transcriptional responses that become apparent in its absence.

GO analysis revealed distinct functional enrichments between these groups: WT-responsive genes were associated with hormone signaling and abiotic response pathways, whereas buffered genes were enriched for developmental processes and responses to metal ions (Fig. 5; Supplementary Data S6). These results suggest that OsSLR1 regulates multiple biological processes through distinct transcriptional modes.

**Figure 5.**
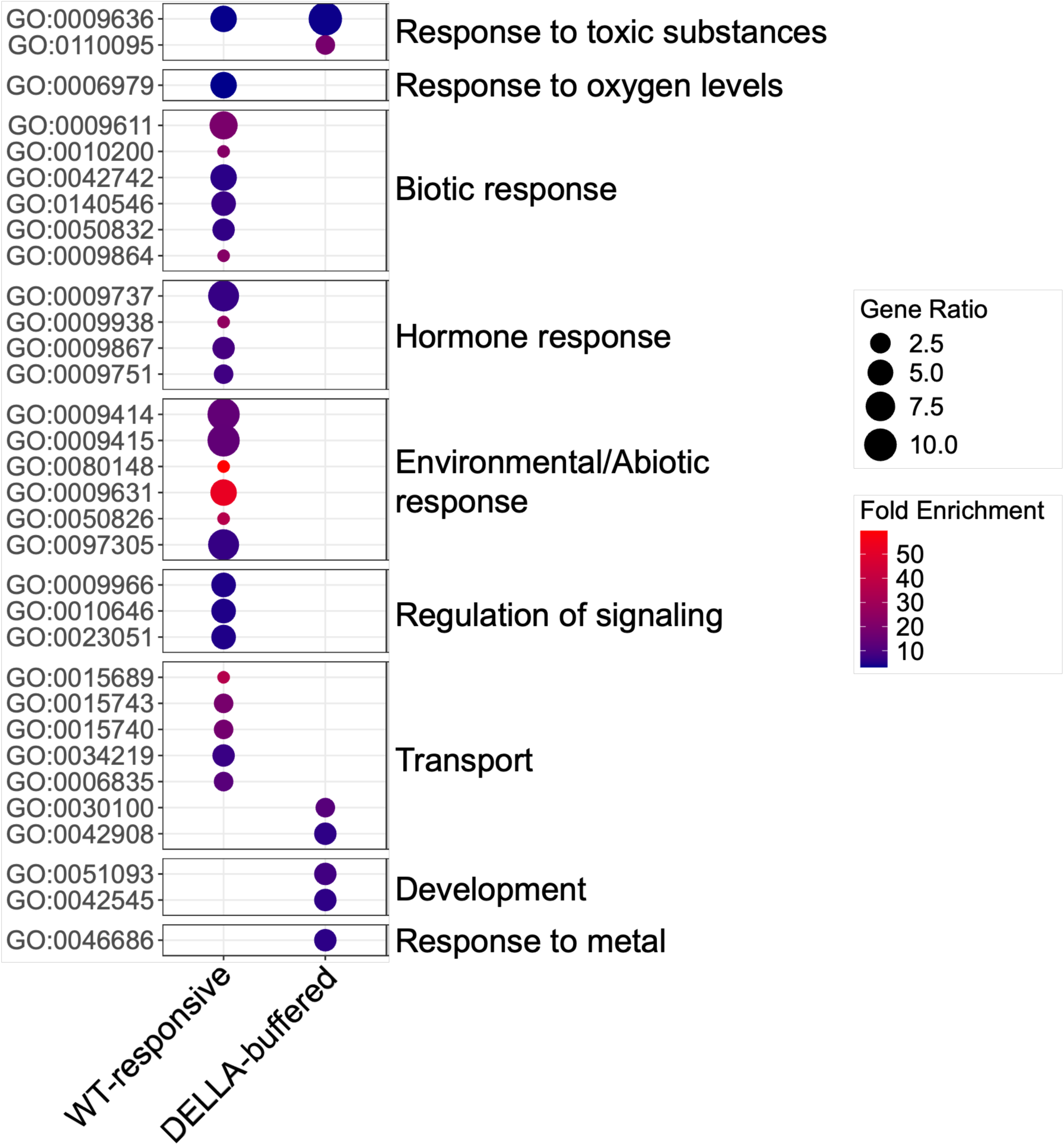
Functional enrichment of OsSLR1-dependent gene expression categories in *O. sativa* roots. Dot plot showing GO biological processes enriched among PEG-responsive DEGs classified as WT-responsive or buffered. Dot size represents gene number, and color intensity indicates enrichment level. Distinct functional categories are associated with each group. Labels on the right correspond to author-assigned functional groupings as in Fig. 3; full GO identifiers and term names are listed in Supplementary Data S6. **Alt text:** Dot plot of enriched Gene Ontology biological processes for WT –responsive and buffered gene sets in *O. sativa*, with dot size showing gene number and colour showing enrichment, grouped into functional categories.

Analysis of TF enrichment revealed that, in contrast to *A. thaliana*, most TFs were shared between WT-responsive and buffered networks in *O. sativa*, suggesting a higher degree of overlap between regulatory modules (Fig. S5; Supplementary Data S7). Together, these findings indicate that OsSLR1 modulates both activation and constraint of transcriptional responses to PEG –induced water limitation in *O. sativa* roots.

### DELLA-mediated transcriptional networks show partial conservation and divergence between *A. thaliana* and *O. sativa*

To assess the conservation of DELLA-dependent responses between species, we compared transcriptional changes among orthologous genes in *A. thaliana* and *O. sativa*. Analysis of fold-change differences across conserved DEGs (i.e. orthologous genes in both species that are either WT-responsive or DELLA –buffered DEGs) revealed largely distinct responses between orthologs (Fig. 6A). Indeed, of the 75 orthogroups represented in Fig. 6A, only six (8%) responded in the same direction in both species across all comparisons, whereas 66 (88%) responded in opposite directions in at least one comparison, and three (4%) showed discrepant behavior among orthogroup members (Supplementary Data S8; see Materials and Methods). This indicates that transcriptional outputs downstream of DELLA signaling have diverged qualitatively within conserved responsive genes between the two species. Nevertheless, GO enrichment analysis of these orthologous DELLA-dependent genes revealed shared functional categories, largely related to metabolism, transport, hormone responses, and responses to environmental stimuli, together with species –specific categories (Supplementary Fig. S6; Supplementary Data S9). Thus, despite the limited conservation of individual transcriptional responses, some functional categories are shared between the two species.

**Figure 6.**
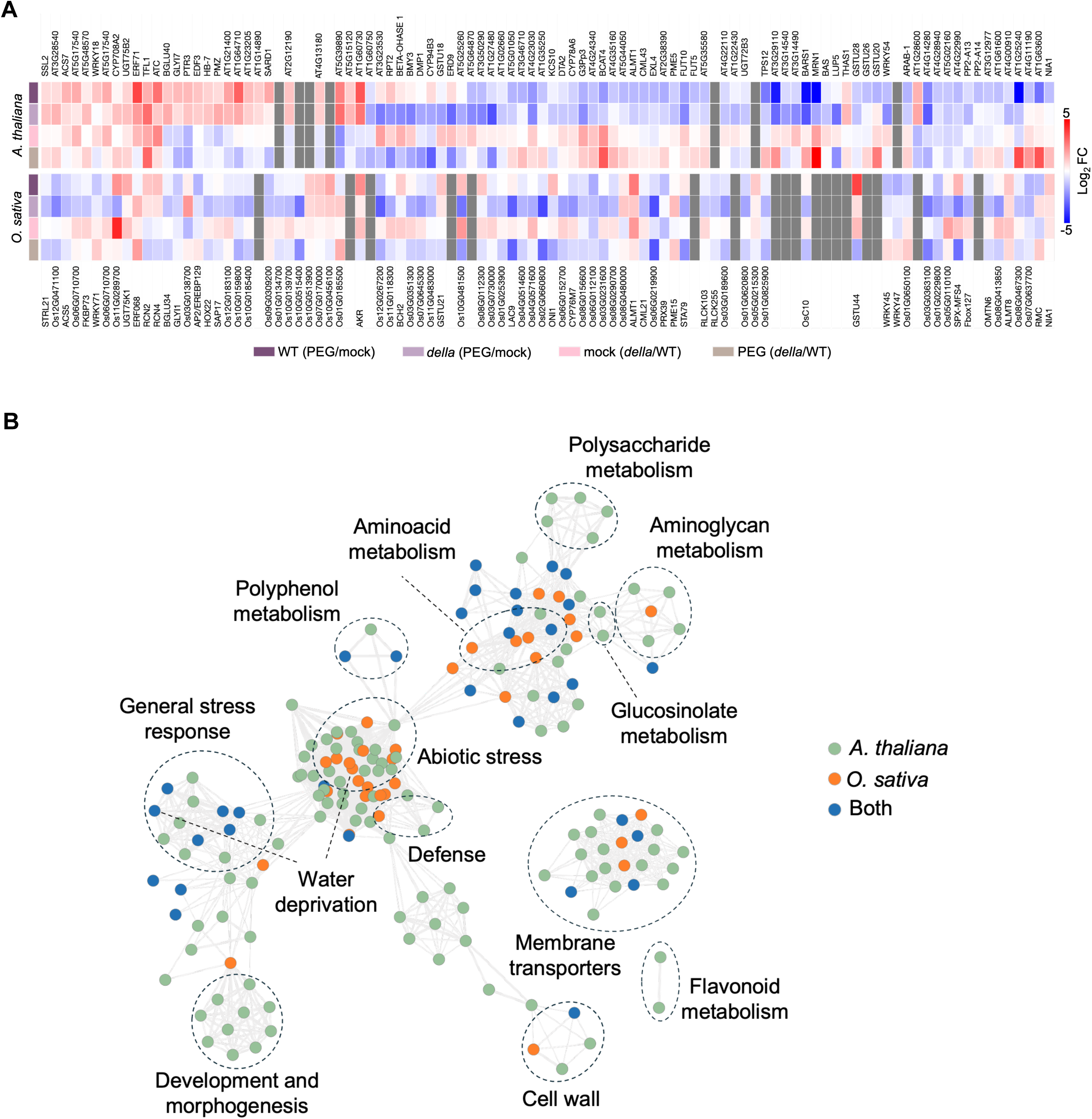
Conservation and divergence of DELLA-associated transcriptional responses between *A. thaliana* and *O. sativa*. **A)** Heatmap of log_2_ fold changes for orthologous genes that are differentially expressed in response to PEG in a DELLA-dependent manner in both species. Genes are labeled with their gene IDs. Because ortholog relationships are frequently many-to-many, an orthogroup may contain different numbers of genes in each species; grey indicates positions without a corresponding ortholog within an otherwise shared orthogroup (see Materials and Methods). Rows were ordered manually to group orthogroups with related expression patterns. **B)** Comparison of GO biological processes enriched among DELLA-dependent DEGs in both species. GO terms were summarized with REVIGO and visualized as a semantic-similarity network in Cytoscape. Each dot represents an enriched term; colors indicate terms shared between species (blue), or specific to *A. thaliana* (green), or *O. sativa* (orange). Edges represent semantic similarity between GO terms, calculated using the SimRel metric, and do not represent biological interactions. All edges present in the REVIGO Cytoscape output were retained, with no additional edge-weight threshold applied in Cytoscape. **Alt text:** Heatmap of fold changes for orthologous DELLA-dependent genes in *A. thaliana* and *O. sativa* labelled with gene identifiers, and a network of enriched Gene Ontology terms coloured by whether they are shared or species –specific.

Then, we examined the GO enrichment across the full set of DELLA –dependent genes in both species. Several general biological processes were enriched in both *A. thaliana* and *O. sativa*, including “response to water deprivation”, suggesting the presence of a shared core response to PEG –induced water limitation (Fig. 6B and Fig. S7; Supplementary Data S10). However, species-specific enrichments were also observed, including development and morphogenesis, and flavonoid metabolism in *A. thaliana* (Fig. 6B and Fig. S7), indicating lineage-specific specialization. We note that the apparent absence of enrichment of a given category in one species should not be interpreted as evidence of biological absence, since gene-set size, statistical power and annotation completeness differ between the two species (see Discussion).

Lastly, we inferred putative DELLA-associated transcriptional regulatory networks based on TF binding site enrichment in both species. Thirteen TFs were conserved between the *A. thaliana* and *O. sativa* networks, which represent a substantial fraction of the *O. sativa* network (Fig. 7A and B). Nonetheless, most TFs were species-specific (Fig. 7A and B), and analysis of downstream target gene expressions revealed substantial divergence in qualitative regulatory relationships downstream of the 13 conserved orthologs (belonging to 11 orthogroups) (Fig. 7C; Supplementary Data S11).

**Figure 7.**
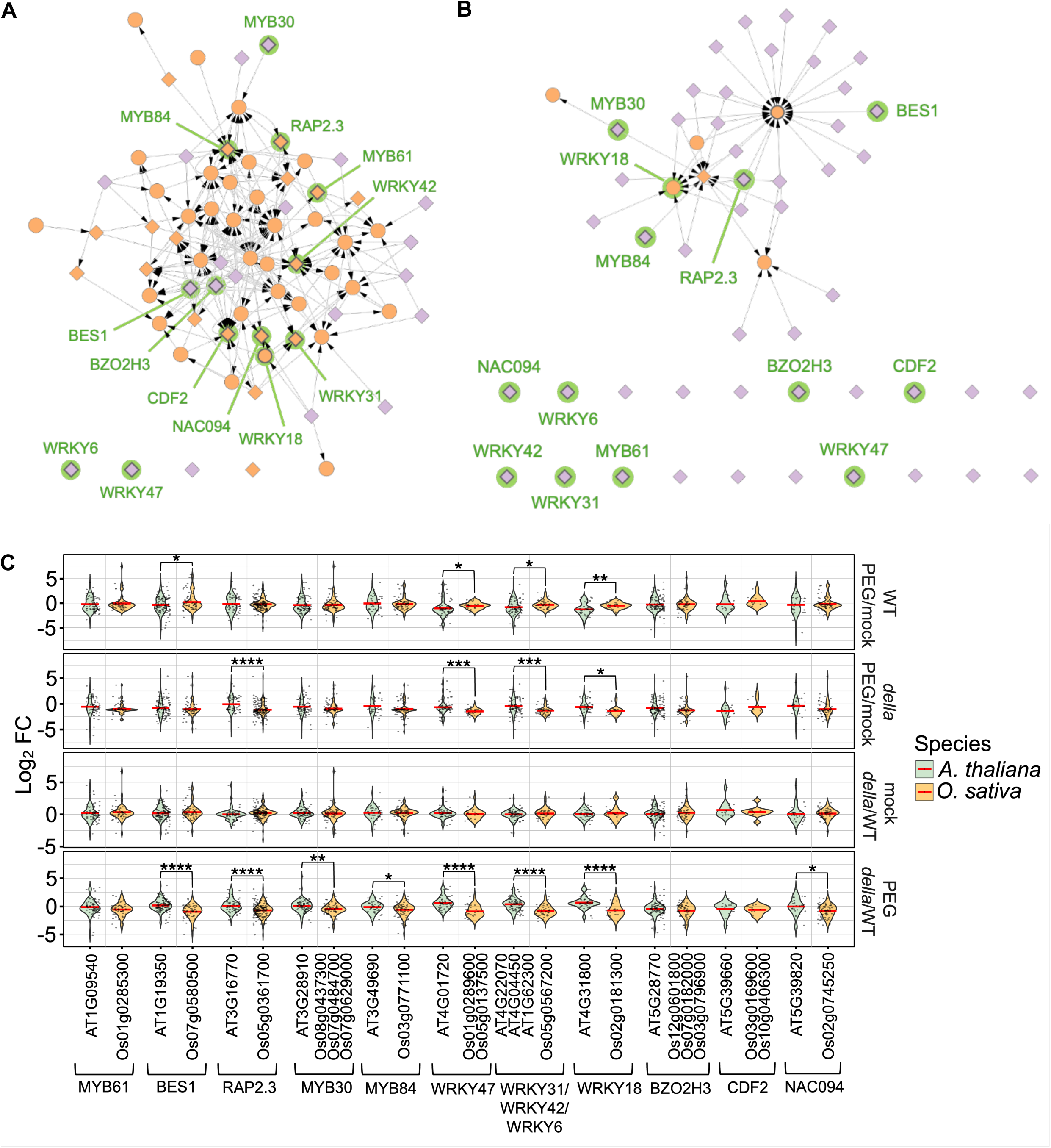
Inferred transcription factor associations within DELLA-dependent gene sets in *A. thaliana* and *O. sativa*. **A)** Inferred transcription factor (TF) associations with DELLA-dependent DEGs in *A. thaliana*, based on enrichment of TF binding sites. **B)** Equivalent analysis in *O. sativa*. In both panels, TFs with enriched binding sites are shown in violet, TFs that are also differentially expressed are shown in orange, and TFs reported as DELLA interactors are indicated by diamonds. TFs with orthologs present in both species are highlighted in green. **C)** Distribution of expression changes (log_2_ fold change) of predicted target genes associated with conserved TF ortholog groups across the different transcriptomic comparisons in both species. Statistical analyses were performed using two-way ANOVA followed by post-hoc tests (* p < 0.05; ** p < 0.01; *** p < 0.001; **** p < 0.0001). The depicted associations are inferred from binding-site enrichment and do not establish direct regulatory interactions. **Alt text:** Two transcription factor association networks, one for *A. thaliana* and one for *O. sativa*, with nodes coloured by enrichment and expression status, and violin plots of predicted target gene fold changes for conserved transcription factor groups.

Altogether, these results indicate that DELLAs act as conserved regulators of root responses to PEG-induced water limitation. While the core regulatory pathway is conserved, broader downstream transcriptional programs exhibit both shared and lineage-specific features consistent with evolutionary rewiring. Differences between the experimental systems across the two species may also contribute these divergence patterns (see Discussion).

## Discussion

Our phenotypic data analyses support the established role of DELLA proteins as central regulators of growth responses under conditions of reduced water availability. In both *A. thaliana* and *O. sativa*, loss of DELLA function reduced the sensitivity of root growth and development to PEG-induced water limitation. Although the specific developmental outputs differed between species (primary root elongation in *A. thaliana* and crown root development in *O. sativa*), these results are consistent with a conserved role for DELLAs in modulating growth responses to reduced water potential (Achard *et al.,* 2008; Serrano-Mislata *et al.,* 2025). Moreover, the distinct phenotypic manifestations highlight the flexibility of this regulatory module across different root architectures.

A central observation of our study is the presence of two distinct modes of DELLA-dependent transcriptional responses to water limitation, here described as “WT-responsive” and “buffered”. The WT-responsive category is consistent with the well-established role of DELLAs enabling or modulating stress-induced transcriptional changes. In contrast, the buffered category comprises genes whose responsiveness to reduced water availability becomes apparent primarily in the absence of DELLAs. This pattern is consistent with a role for DELLAs in constraining transcriptional responsiveness, thereby limiting the extent of gene expression changes under stress conditions. Importantly, our data do not directly demonstrate the underlying mechanisms of this effect; rather, they define a class of genes whose behavior is compatible with a buffering or stabilizing role for DELLAs at the transcriptional level. A stronger or exclusive response in the DELLA-deficient background could equally arise from differences between genotypes in basal expression, developmental stage or root cellular composition, from a different perceived severity of the stress, or from secondary consequences of the pleiotropic phenotypes of DELLA-null mutants. In whole-root profiling, a shift in the relative abundance of cell types between genotypes would also produce a comparable signature. Distinguishing between these possibilities will require cell-type-resolved profiling and direct measurements of DELLA occupancy at candidate loci. Such a role could contribute to maintaining coordination between stress responses and developmental programs, preventing excessive or potentially maladaptive transcriptional changes under fluctuating environmental conditions (von Heckel *et al.,* 2016; Lu *et al*., 2023; Kadelka, 2026).

Our comparative analyses support a model (Fig. 8) in which DELLA proteins function as conserved regulatory hubs, while the downstream transcriptional programs they influence have diverged substantially between species (Briones-Moreno *et al.,* 2023). In this context, what is conserved between *A. thaliana* and *O. sativa* is the requirement for DELLA function in adjusting root growth and transcriptional responses to reduced water availability; the developmental output through which this requirement is expressed, the magnitude of the effects and the identity of the responsive genes are largely species-specific. Although a subset of TFs appears to be shared between *A. thaliana* and *O. sativa*, the overall transcriptional responses of their predicted target genes differ markedly, indicating substantial qualitative divergence in regulatory outputs. This aligns with the hypothesis that evolutionary changes in *cis*-regulatory elements or network connectivity, rather than in core signaling components, contribute to species-specific adaptation to water limitation (Artur and Kajala, 2021; Marand *et al*., 2023). Moreover, the predictive power of this analysis is supported by the fact that five of these conserved TFs have been previously characterized as being involved in responses to water availability in *A. thaliana*: AT1G09540 (AtMYB61) (Liang *et al.,* 2005), AT1G19350 (BES1) (Korwin Krukowski *et al.,* 2022), AT3G16770 (RAP2.3) (Papdi *et al*., 2015), AT3G28910 (AtMYB30) (Zhang *et al.,* 2023) and AT4G31800 (AtWRKY18) (Chen *et al*., 2010).

**Figure 8.**
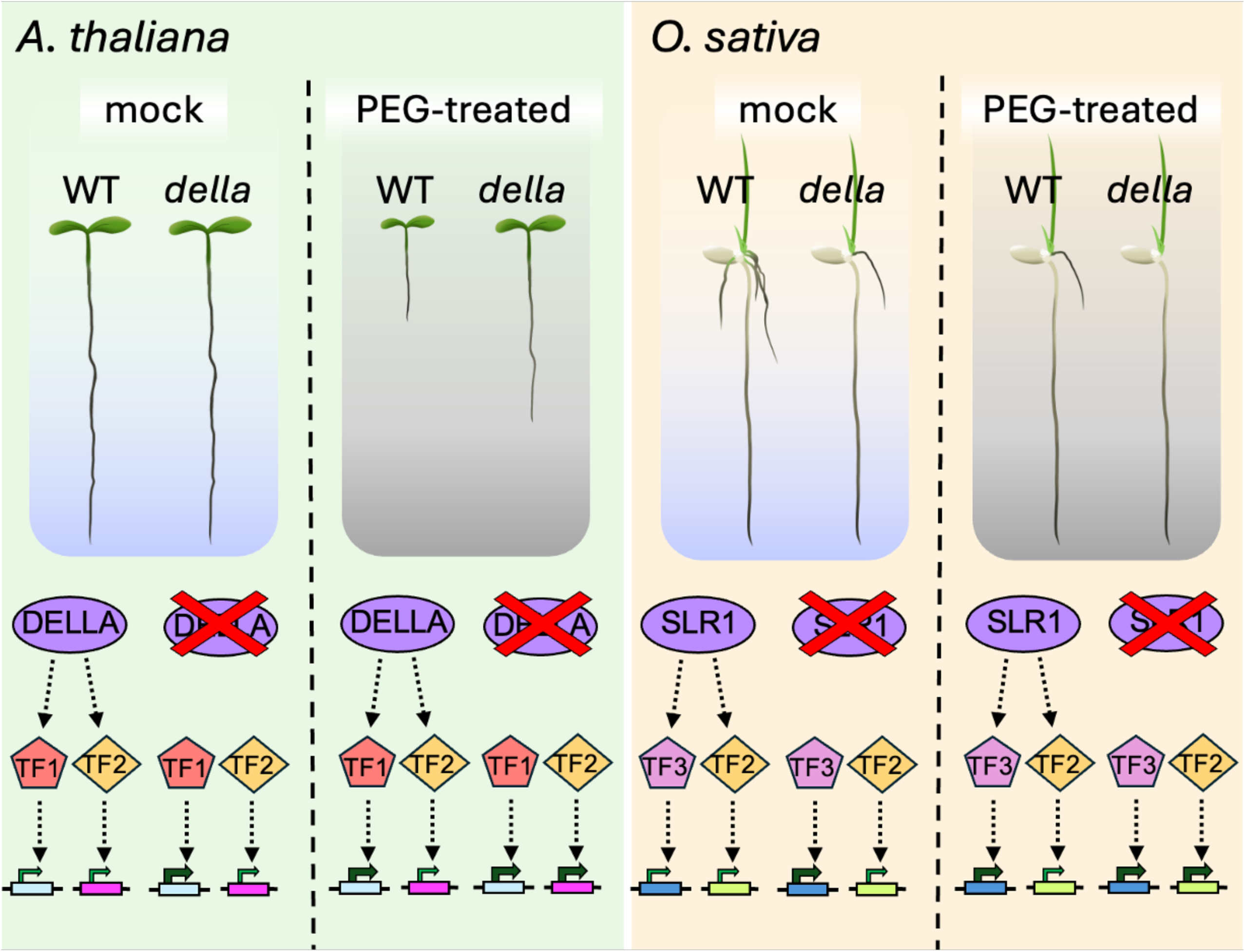
Model summarizing DELLA-associated transcriptional responses to water limitation in roots. Schematic representation of the proposed role of DELLA proteins in modulating root responses to PEG-induced water limitation in *A. thaliana* and *O. sativa*. DELLA proteins are associated with both the regulation of PEG-responsive genes and the apparent constraint of additional transcriptional responses that become evident in their absence. A subset of transcription factors is conserved between species, whereas downstream transcriptional responses differ. Dashed arrows denote inferred DELLA-TF and TF-target associations, which have not been experimentally demonstrated in this study, and green arrow thickness represents relative gene expression levels. **Alt text:** Schematic model showing *A. thaliana* and *O. sativa* seedlings under mock and PEG conditions in wild type and DELLA mutants, with dashed arrows from DELLA to transcription factors to target genes and bars indicating expression levels.

When interpreting these cross-species differences, it is also important to consider that the two experimental systems differ in several respects beyond the genetic perturbation, including root system architecture and root types, developmental stage, growth medium, photoperiod, temperature and treatment duration. The same nominal PEG8000 concentration should therefore not be assumed to impose an equivalent effective stress in both species, and the different apparent PEG sensitivities of *A. thaliana* and *O. sativa* roots may partly reflect these experimental differences. Consequently, shared responses provide the most direct basis for cross-species comparison, whereas species-specific responses should be interpreted with this limitation in mind. Similarly, differences in genome annotation completeness and in the coverage of transcription factor binding-site models between the two species can influence the number of enriched GO terms and the size and connectivity of the inferred association networks; we therefore do not interpret such differences as evidence of differences in regulatory complexity between species. For these reasons, we present evolutionary rewiring of DELLA-associated networks as one plausible contributor to the observed divergence rather than as an established explanation.

An important consideration in this study is the use of PEG-induced water limitation as an experimental system. While PEG-based approaches do not fully recapitulate the complexity of soil drought, they provide a controlled framework to impose defined reductions in water potential, allowing the dissection of transcriptional responses to osmotic and hydraulic components of stress (Gonzalez *et al.,* 2024). Therefore, our findings should be interpreted as reflecting responses to reduced water potential under controlled conditions, rather than encompassing the full spectrum of drought responses occurring in natural environments. We also note that the similar primary root elongation of wild type and At*dellaKO* seedlings at 25% PEG8000 does not guarantee an equivalent developmental stage at the time of sampling, and this remains a possible contributor to the transcriptional differences reported here. In addition, our transcriptomic analyses were performed on whole-root samples, which likely integrate signals from multiple cell types and developmental zones (Illouz-Eliaz *et al*., 2025). As a result, cell type-specific or spatially restricted transcriptional responses may be masked, and the apparent patterns of responsiveness and constraint described here should be understood as emerging from the composite behavior of heterogeneous root tissues.

Despite these limitations, our findings provide a framework for understanding how DELLA proteins contribute to the organization of root transcriptional responses under reduced water availability. By distinguishing between responsive and constrained modes of gene expression, this study highlights an additional layer of regulation that may contribute to the coordination of growth and stress responses. Elucidating how these modes are implemented and how they vary across species may offer new opportunities to refine strategies for improving plant performance under water-limited conditions.

## Supplementary data

The following supplementary data are available at JXB online.

**Fig. S1.** Effect of PEG-induced water limitation on crown root formation in *O. sativa*.

**Fig. S2.** Principal component analyses of RNA-seq replicates in *A. thaliana* and *O. sativa*.

**Fig. S3.** Overlap of DELLA-dependent DEGs across conditions in *A. thaliana* and *O. sativa*.

**Fig. S4.** Inferred TF associations for DELLA-dependent gene sets in *A. thaliana*.

**Fig. S5.** Inferred TF associations for DELLA-dependent gene sets in *O. sativa*.

**Fig. S6.** Functional categorization of PEG-responsive and DELLA-dependent *A. thaliana* and *O. sativa* orthologs.

**Fig. S7.** Functional categorization of DELLA-dependent gene expression across species.

**Data S1.** Raw root phenotypic measurements, sample sizes and statistical analyses for *A. thaliana* and *O. sativa*, including representative images of *A. thaliana* seedlings.

**Data S2.** Differential expression and genotype-by-environment (G×E) analyses of root transcriptomes under PEG-induced water limitation in *A. thaliana* and *O. sativa*.

**Data S3.** Numbers and identities of up –and downregulated genes within each differentially expressed gene set in *A. thaliana* and *O. sativa*.

**Data S4.** Gene lists corresponding to hierarchical clustering of DELLA –dependent transcriptional responses in *A. thaliana* and *O. sativa*.

**Data S5.** Genes showing significant genotype-by-environment (G×E) interactions classified as WT-responsive or buffered in *A. thaliana* and *O. sativa*.

**Data S6.** Gene Ontology enrichment analyses of WT-responsive and buffered gene sets in *A. thaliana* and *O. sativa*.

**Data S7.** TF enrichment and inferred associations for WT-responsive and buffered gene sets in *A. thaliana* and *O. sativa*.

**Data S8.** Expression values (log_2_ fold change), gene identifiers and gene symbols of orthologous DELLA-dependent genes, together with the classification of orthogroups by direction of response, in *A. thaliana* and *O. sativa*.

**Data S9.** Gene Ontology enrichment analyses of the orthologous DELLA –dependent genes shown in Fig. 6A.

**Data S10.** Gene Ontology terms enriched among DELLA-dependent genes shared between or specific to *A. thaliana* and *O. sativa*.

**Data S11.** Predicted TF–target relationships and expression values for conserved transcription factor ortholog groups.

## Acknowledgements

We would like to thank Ludovico Dreni and Fernando Andrés for helping with rice husbandry in the early stages of the project. We are grateful to Javier Forment for critical help with the initial transcriptomic bioinformatic pipeline. We are also thankful for all the input and support from all the members of the Plasticity Lab.

## Author contributions

C.W., M.A.B and B.C. designed the research. C.W. and B.C. performed all the experiments. C.W., M.A.B. and B.C. analyzed the data. C.W. created the figures. D.A., M.A.B. and B.C. provided project guidance and supervision. All authors discussed the data. B.C. wrote the initial draft, and all authors reviewed and edited the manuscript.

## Data availability

The RNA-sequencing data and corresponding metadata are deposited at the National Center for Biotechnology Information (NCBI), BioProject GSE328143. All relevant data are available from the corresponding author upon request. There are no restrictions on data availability.

## Abbreviations

DEG: Differentially Expressed Gene
GA: Gibberellin
GO: Gene Ontology
GxE: Genotype-by-Environment
L*er*: Landsberg *erecta*
MS: Murashige and Skoog
PEG: Polyethylene Glycol
SLR1: SLENDER1
TF: Transcription Factor
TPM: Transcripts per million
WT: Wild Type

